# Analysis of the progression of cervical cancer in Guatemala- from pre-malignancy to invasive disease

**DOI:** 10.1101/2024.10.25.620123

**Authors:** Emma Robinson, Isabel Rodriguez, Victor Argueta, Yi Xie, Hong Lou, Rose Milano, Hyo Jung Lee, Laurie Burdett, Sambit K. Mishra, Meredith Yeager, Lisa Mirabello, Michael Dean, Roberto Orozco

**Affiliations:** HLA Immunogenetics, Basic Science Program, Frederick National Laboratory for Cancer Research, Gaithersburg, MD USA; Laboratory of Translational Genomics, Division of Cancer Epidemiology and Genetics, National Cancer Institute, Gaithersburg, MD USA; Hospital General San Juan de Dios, Guatemala City, Guatemala; Cancer Genetics Research Laboratory, Division of Cancer Epidemiology and Genetics, Frederick National Laboratory for Cancer Research, Gaithersburg, MD USA

**Keywords:** cervical cancer, precancer, gene expression, tumor microenvironment, human papillomavirus, cancer progression, immune surveillance

## Abstract

To better understand cervical cancer progression, we analyzed RNA from 262 biopsies from women referred for colposcopy We determined HPV type and analyzed the expression of 51 genes. HPV31 was significantly more prevalent in precancer than stage 1 cancer and invasive cancer (p < 0.0001) and HPV16 increased in invasive disease (p < 0.0001). *CCNE1, MELTF*, and *ULBP2* were significantly increased in HPV16-positive compared to HPV31 precancers while *NECTIN2* and *HLA-E* expression decreased. Markers of the innate immune system, DNA repair genes, and cell cycle genes are significantly increased during cancer progression (p = 0.0001). In contrast, the *TP53* and *RB1* tumor suppressor gene expression is significantly decreased in cancer cells. TheT cell markers *CD28* and *FLT3LG* expression decreased in cancer while *FOXP3, IDO1*, and *ULBP2* expression increased. There is a significantly higher survival rate in individuals with increased expression of *CD28* (p = 0.0005), *FOXP3* (p = 0.0002), *IDO1* (p = 0.038), *FLT3LG* (p = 0.026), *APOBEC3B* (p = 0.0011), and *RUNX3* (p = 0.019), and a significantly lower survival rate in individuals with increased expression of *ULBP2* (p = 0.035). These results will help us understand the molecular factors influencing the progression of cervical precancer to cancer.

## 1. Introduction

A total of 15 human papillomavirus types are recognized as oncogenic/high risk (hr) for cervical cancer. Over 80% of women will acquire an infection from at least one of these hr-HPV types in their lifetime, but over 90% will clear the infection in 1-2 years [1]. About 5% of women with a persistent hr-HPV infection will progress to cervical precancer, and an even smaller percentage will progress to localized and then to invasive disease [2]. Of the hr-HPV types, HPV16, HPV18, and HPV45 are often the three most frequent types found in invasive cancers and are more oncogenic than other hrHPV types [3]. However, the distribution of the 15 hrHPV types varies considerably by geographic location [4].

Of the estimated 604,000 annual global cases of cervical cancer and 342,000 deaths due to cervical cancer in 2020 [5], 595,115 (91%) of cancer and 319,713 (92%) deaths occurred in low and middle-income countries (LMICs) ([6]). Therefore, it is critical to study the prevalence of HPV types from precancer to cancer lesions to understand the molecular changes occurring during cancer progression in understudied regions of the world.

The availability of cervical tissue from cancer-free, precancer, and cancer cases allows the study of the transition from HPV infection to precancer, localized cancer, and invasive cancer. Previously, a study of 128 US samples identified increases in the expression of genes involved in cell proliferation and DNA repair during the progression to cancer and a decrease in the estrogen receptor gene expression [7] Another study of 28 US samples identified *HOXC10* expression as a marker of invasiveness [8].To date, there are few studies of cervical cancer progression in LMIC populations.

The strongest predictor of the progression of HPV infection or cervical precancer to cancer is HPV type. Analysis of the rate of progression to cancer has shown that HPV16, HPV18, and HPV45 have the highest rates. Many studies demonstrate that HPV16, 18, and 45 are the most frequent HPV types in advanced/invasive cervical cancer, whereas these types are often not the dominant hrHPV types in individual populations. [4, 7].

The molecular basis of the differences in progression remains poorly understood. Therefore, we studied gene expression and HPV types in 262 samples of cancer-free controls, precancer, and local cancer in the same hospital in Guatemala and compared this data with 454 invasive cancer samples from the same city. We identified several DNA repair, cell cycle, and immunological genes associated with the progression from controls to precancer to cancer and cancer survival.

## 2. Materials and Methods

### 2.1 Patient samples and histology

Formalin-fixed paraffin-embedded (FFPE) tissue samples were selected from women referred for colposcopy to evaluate cervical lesions at the Hospital General San Juan de Dios. The Ministry of Health of Guatemala approved the study, and samples were coded. Slides from each sample were examined by a pathologist, and detailed histology was recorded. Samples were grouped into **Control** (low-grade precancer and cancer-free), **Precancer** (high-grade precancer), and **Cancer** (stage 1). Cancer was further subdivided into histological subtypes (adenocarcinoma, squamous cell carcinoma). Data on advanced, invasive cervical cancers came from the Instituto Nacional de Cancerologia (INCan) in Guatemala and have been described previously [9]. Tissue blocks were placed on a microtome, and three 10um sections were collected, cleaning the instrument and blade between samples.

### 2.2 RNA extraction

Total RNA was extracted from FFPE cervical tissue samples obtained from 262 women referred for colposcopy in a single hospital in Guatemala. The FFPE to Pure RNA kit with Ionic® Purification system (Purigen Biosystems, Pleasanton, CA) was utilized for extraction, following the manufacturer’s protocol. FFPE tumor scrolls were lysed with reagents in the RNA FFPE kit, and samples were loaded onto Purigen cartridges. The extracted RNA was quantitated using Qubit RNA Broad Range Assay (ThermoFisher, Waltham, MA).

### 2.3 Gene expression analysis

The gene expression assay was conducted on NanoString’s nCounter MAX/FLEX system using a custom nCounter XT Codeset (NanoString Technologies, Seattle, WA). The custom Codeset contained 77 probes covering 51 genes and probes for HPV16, HPV18, and HPV45, as well as negative and positive controls (see **Supplementary Table S1** for gene and probe information). The custom Codeset included four housekeeping genes (*GAPDH, G6PD, TBP*, and *ACTB*) and the manufacturer’s six positive and eight negative controls.

For the assay, 250 ng of total RNA for each sample was normalized to 5ul. The hybridization of the sample to the custom nCounter XT Codeset was performed according to the manufacturer’s protocol for nCounter XT CodeSet Gene Expression Assays using the ProFlex PCR System (ThermoFisher, Waltham, MA) at 65°C for 18 h. After hybridization, samples were loaded on cartridges using the Nanostring nCounter Prep Station 5s, and the cartridges were scanned on the nCounter Digital Analyzer according to the manufacturer’s protocol. Per the manufacturer’s recommendations, nSolver Analysis Software 4.0 (NanoString Technologies, Seattle, WA) was used for quality control assessment and data normalization. Expression levels of target genes were normalized to positive and negative controls and housekeeping genes.

### 2.4 Analysis of gene expression and cancer subtype

The dataset used in this analysis contained 219 unique patient samples, and the variables created for this analysis were age category and histological category (**Supplementary Fig. S1**). The histological category consisted of Controls, Precancer, and Stage 1 Cancer, and the age category divided the subjects into women fifty and younger (assumed to be premenopausal) and above fifty (menopausal). The descriptive statistics of the non-gene variables are shown in **Tables 1** and **2**. In total, 40 samples had expression measured twice, and two were measured in triplicate. For each set of duplicate observations, we used the maximum value for each gene. Of the original 262 samples, 43 were dropped from the analysis due to poor QC measures (housekeeping gene normalization flags) and low overall gene expression.

**Table 1.** Descriptive Statistics.

| Histology (219) | Frequency (Percent) |
| --- | --- |
| Adenocarcinoma | 2 (1%) |
| Squamous Cell Carcinoma | 88 (40%) |
| High-Grade Precancer | 65 (29%) |
| Low-Grade Precancer | 13 (6%) |
| Control | 51 (23%) |
| HPV Type (154) | Frequency (Percent) |
| HPV16 | 43 (28%) |
| HPV18 | 12 (8%) |
| HPV31 | 37 (24%) |
| HPV33 | 11 (7%) |
| HPV35 | 1 (1%) |
| HPV39 | 2 (1%) |
| HPV45 | 11 (7%) |
| HPV51 | 3 (2%) |
| HPV52 | 6 (4%) |
| HPV56 | 5 (3%) |
| HPV58 | 20 (13%) |
| HPV59 | 3 (2%) |
| Age Category (186) | Frequency (Percent) |
| Premenopausal ($\leq 50$) | 140 (75%) |
| Post-menopausal ($> 50$) | 46 (25%) |
| Histologic Category (219) | Frequency (Percent) |
| Control | 64 (29%) |
| Precancer | 65 (30%) |
| Cancer | 90 (41%) |

The normality of the expression data for each gene was assessed, and all but two of the genes were not normally distributed. Transformations were conducted to normalize the data. Logarithmic transformations were attempted for each gene, and 28 genes were successfully normalized. The distributions of the genes that could not be normalized through logarithmic transformations had their high outliers clipped, successfully normalizing fifteen more genes. The five remaining genes (*ASCL1, CCNA1, CD94, KIR*, and *CD96*) could not be normalized through clipping or logarithmic transformations. These genes were analyzed using nonparametric tests. *LHX8* was excluded from the analysis because no individuals showed expression of *LHX8*.

One-way ANOVA tests and pairwise t-tests were conducted to analyze the genes that had been successfully normalized. The age category was analyzed as a potential interactor in these analyses. Stratified results were presented for genes in which age category and histological category were found to interact. Nonparametric Kruskal-Wallis and Dwass-Steel-Critchlow-Fligner (DSCF) pairwise tests were conducted to analyze the genes that had not been successfully normalized. Age was not analyzed as a potential interactor in these analyses due to the lack of suitable nonparametric interaction tests. The significance level was set as α = 0.05, and a Bonferroni correction was applied to pairwise analyses. Genes significantly increased or decreased in cancer samples were then examined in the TCGA cohort to determine the association with survival. SAS 9.4 and R 4.3.1 were used to conduct this analysis.

The Nanostring assay was also performed on eighteen cervical cancer cell lines, as was a transcriptome sequencing assay. A correlation analysis was conducted to compare the results of the two assays. Gene expression levels measured by Nanostring were very strongly (r > 0.90) correlated with gene expression levels measured by Oxford Nanopore transcriptome sequencing [10] in the cell lines ME-180, MS751, SNU-1245, SNU-1299, SNU-17, and SNU-682. Gene expression levels measured by Nanostring were strongly (0.70 < r ≤ 0.90) correlated with gene expression levels measured by transcriptome sequencing in the C4-I, HT-3, SCC154, SNU-1000, SNU-1005, SNU-487, SNU-703, SNU-778, SNU-902, SW756, and SiHa cell lines. Gene expression levels measured by Nanostring were moderately (0.51 < r ≤ 0.70) correlated with gene expression levels measured by transcriptome sequencing in the HeLa cell line (**Supplementary Table S2**). These findings suggest that the Nanostring assay has equivalent validity to the Oxford Nanopore transcriptome sequencing assay.

### 2.5 HPV sequencing and phylogenetics

Purified FFPE DNA samples were used for a targeted sequencing panel (NCI CHANGeS) (National Cancer Institute Carcinogenic HPV All Next Generation Sequencing) amplifying overlapping segments of the entire viral genome for 13 HPV types using Ampliseq technology from Ion Torrent and are an extension of this method [11]. HPV16, 18, and 45 data were validated using the nanostring probes for those types. Samples were initially classified by their dominant HPV type, as many samples were positive for multiple HPV types at low levels. Samples with high reads for multiple HPV types were classified as multitype. Invasive cancer samples were HPV typed by either GP-5+/6+ PCR amplification and Sanger sequencing (316 samples) or the CHANGeS assay (138 samples). Chi-squared tests were conducted to investigate the difference in the proportion of HPV types in precancer, stage 1, and invasive cancer. T-tests were conducted to examine differential gene expression between HPV16-positive and HPV31-positive precancer samples. A consensus viral sequence was generated, and HPV16 and HPV31 samples were aligned. Samples with a complete HPV type alignment were included, regardless of how they had been classified previously.

Phylogenetic trees of the consensus HPV sequences were produced in MEGA along with controls for viral sublineages to assign sublineage [11]. Chi-squared tests were conducted to examine the change in the proportion of different HPV16 sublineages between precancer and stage 1 cancer.

### 2.6 Sanger sequencing

Primers were designed using Primer3 [12] to amplify specific regions of HPV to determine lineage and sublineage (**Supplementary Table S3**) in a subset of samples. Purified PCR products were sequenced on an Applied Biosystems® 3500xL Genetic Analyzer (Thermofisher Scientific). We analyzed the sequences using DNASTAR SeqMan Ultra and BLAST (https://blast.ncbi.nlm.nih.gov/Blast.cgi) to validate HPV type and sublineage further.

### 2.7 Survival analyses

Gene expression and survival data from The Cancer Genome Atlas (TCGA) [13] was downloaded from cbioportal (https://www.cbioportal.org/) [14]. Kaplan-Meier curves were constructed to investigate the relationship between gene expression and survival in individuals with cervical cancer.

## 3.0 Results

### 3.1 Patient samples and HPV typing

To study the transition of cervical cells from infection to precancer and cancer, we analyzed a serial sample of formalin-fixed tissue blocks from 219 women referred for colposcopy. The sample consisted of 64 controls with no lesion or low-grade abnormalities (controls), 65 high-grade precancers, and 90 localized cancers (Stage 1). Nearly all localized cancers were squamous cell carcinoma (SCC) (88/90 92%). A total of 75% of the subjects had their biopsy before age 50 and were considered premenopausal (**Table 1, Supplementary Fig. S1**).

DNA from the tissue blocks was used to amplify and sequence the genome of 13 hr-HPV types to determine the distribution of HPV types in the sample. HPV typing was obtained from 56 precancer and 46 cancer samples, and overall, HPV16, HPV31, and HPV58 were the three most prevalent HPV types (**Table 2**). However, when samples were divided into precancer and localized cancer, the most prevalent type in precancers were HPV31 (32%), HPV16 (23%), and HPV58 (16%), whereas in localized cancers, they were HPV16 (31%), HPV33 (12%), and HPV45 (9%). To understand if this distribution further changes in patients with invasive cancer, we compared our data to that of 454 grade 2-4 cancers obtained from a hospital in the same city and medical system. In invasive cancers, the three most prevalent types are HPV16 (57%), HPV18 (17%) and HPV45 (11%) (**Table 2**).

**Table 2.** Significant results of trend test for the proportion of HPV type during progression.

|  |  |
| --- | --- |
| HPV16 | $p < 0.0001$ |
| HPV18 | $p = 0.0075$ |
| HPV31 | $p < 0.0001$ |
| HPV33 | $p < 0.0001$ |
| HPV58 | $p < 0.0025$ |

HPV types that significantly differ in frequency using a Cochran-Armitage trend test are shown above.

Frequencies and proportions of the histology, HPV type, and age of the samples used in this analysis are shown. Histology is condensed further into the variable Histologic Category, which combines adenocarcinomas with squamous cell carcinomas and low-grade precancers with controls.

### 3.2 Probability of progression to invasive cancer

Examining the distributions of different HPV types showed that certain HPV types are more frequent in cancer than precancer, indicating more rapid progression (**Supplementary Table S4**). To validate this finding, Cochran-Armitage trend tests were conducted to determine if a significant trend in the proportion of HPV types between precancer, stage 1 cancer, and invasive cancer existed (**Fig. 1, Table 2, Supplementary Table S4)**. Five HPV types had significant trends: HPV16 (p < 0.0001), HPV18 (p = 0.00745), HPV31 (p < 0.0001, HPV33 (p < 0.0001), and HPV58 (p = 0.0.0025). The higher frequencies of HPV16 and HPV18 in invasive cancer indicate that these HPV types are more rapidly progressing. In contrast, the higher frequencies of HPV31 and HPV58 in precancer and the higher frequency of HPV33 in precancer and stage 1 cancer indicate that they are more slowly progressing.

**Fig. 1.**
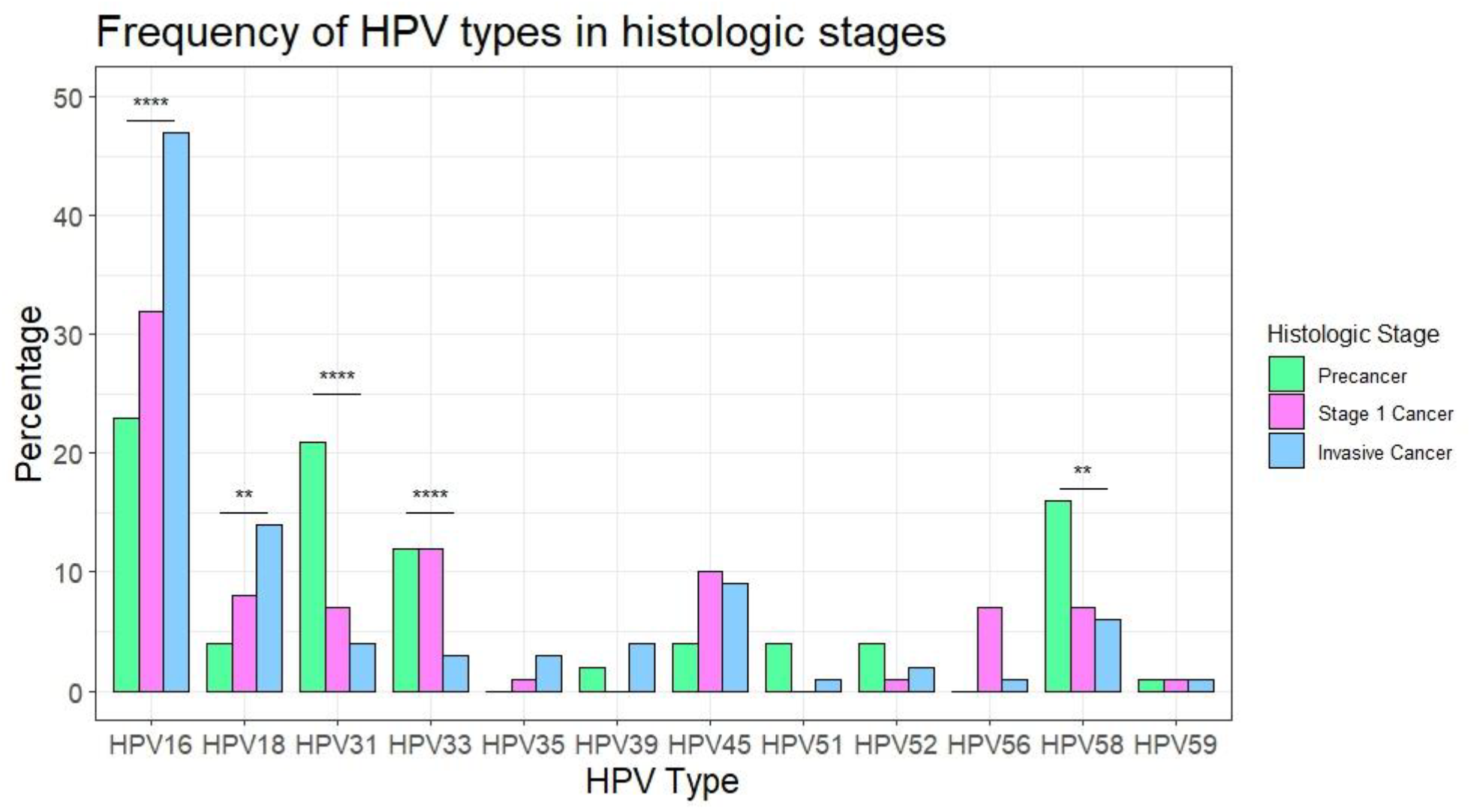
Distribution of HPV type during cervical cancer progression. The percentage of samples in precancer, stage 1 cancer, and invasive cancer are shown by HPV type. HPV68 is included in the panel but was not detected in any precancer or stage 1 cancer therefore it is not shown on this graph.

### 3.3 Comparison of gene expression in HPV16 versus HPV31 precancer

HPV16 and HPV31 are the most prevalent types and show a significant difference in progression from precancer to invasive cancer. HPV16 is a fast-progressing HPV type, while HPV31 is slow-progressing; however, they are closely related evolutionarily. To investigate drivers of cervical cancer progression, we identified immune and cell proliferation genes differentially expressed between precancers with HPV16 and HPV31 (**Fig. 2**). The cyclin E1 gene (*CCNE1*) was significantly increased (p = 0.0081) in HPV16-positive precancers compared to HPV31 positive precancers. CCNE1 can activate cyclin-dependent kinase 2 (CDK2), accelerating the transition through G1/S and G2/M and promoting cell growth. Melanotransferrin (*MELTF*) was also significantly increased in HPV16-positive precancers (p = 0.0082) and not HPV31-positive precancers. MELTF has been implicated in cell proliferation and disease progression in multiple cancers [15-17]. Higher levels of the cyclin genes and melanotransferrin, all of which foster cell growth in HPV16 precancers, are consistent with a higher risk of progression for HPV16 precancers.

**Fig. 2.**
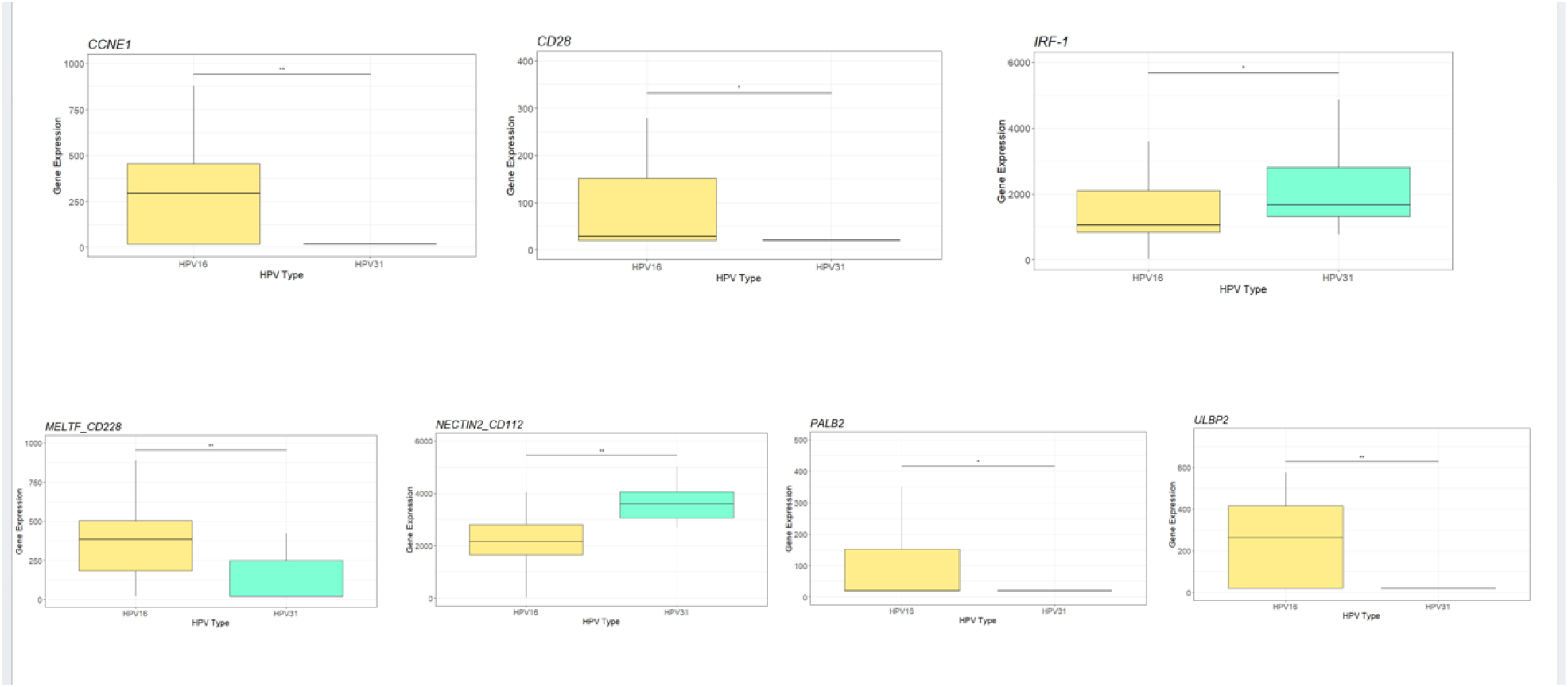
Analysis of gene expression differences in HPV16 and HPV31-positive precancers. These box and whisker plots represent the expression level of various genes in HPV16 and HPV31-positive precancers. Gene expression is shown on the y-axis, while HPV type is on the x-axis. The expression of cell proliferation promoters *CCNE1* and *MELTF*, the NK-cell regulators *ULBP2*, and the DNA-repair gene *PALB2* are increased in HPV16-positive precancers. The expression of the NK-cell co-stimulator and co-inhibitor *NECTIN2*, the immune gene *IRF-1*, and the NK-cell educator *HLA-E* is decreased in HPV16-positive precancers. *** indicates significance of p < 0.001, ** indicates significance of p < 0.01, and * indicates significance of p < 0.05.

Three NK cell ligands, UL16 binding protein 2 (*ULBP2*), nectin cell adhesion molecule 2 (*NECTIN2*), and human leukocyte antigen E (*HLA-E*) showed differential expression between HPV16 and HPV31 precancers. Expression of *ULBP2* is increased (p = 0.0028) in HPV16-positive precancers and not HPV31-positive precancers. Tumor cells secrete soluble ULBP2 to evade NK cells expressing NKG2D [18]. NECTIN2 is either co-stimulatory or co-inhibitory of NK cells, depending on the membrane receptor to which it binds [19]. Expression of *NECTIN2* is decreased (p = 0.0013) in HPV16 precancers and not HPV31 precancers. Expression of *HLA-E* was increased (p = 0.048) in HPV31-positive precancers. *HLA-E* plays a role in the education of NK cells, allowing for an increased immune response [20]. The differential expression of *ULBP2, NECTIN2*, and *HLA-E* indicates that HPV16-positive precancer cells are much more able to evade NK cells than HPV31-positive precancer cells. *CD28* expression is increased in HPV16-positive precancers and the expression of CD28 can stimulate antitumor immunity [21].

Expression of Interferon 1 (*IRF-1*) was decreased in HPV16-positive precancers as compared to HPV31-positive precancers. The role of *IRF-1* in carcinogenesis is unclear and often contradictory, as it is known to inhibit both tumor growth and immune responses [22]. Expression of *PALB2* was increased (p = 0.049) in HPV16-positive precancers as compared to HPV31-positive precancers, indicating a higher rate of DNA-repair in HPV16 precancers.

### 3.4 HPV16 and 31 sublineages and progression

HPV types can be subdivided into lineages and sublineages with known differences in carcinogenicity [23][24]. We performed alignment and phylogenetic analyses of the two most common types of our sample, HPV16 and HPV31. The 78 HPV16 samples with adequate sequence coverage could be divided into A1 (50%), D2 (30%), and D3 (20%) (**Supplementary Table S5**). This finding agrees with previous data from Guatemalan invasive cancers, demonstrating that the HPV16 D2 and D3 sublineages are common in this country. Chi-squared tests were conducted to identify sublineages that were more common in either precancer or precancer. D3-positive samples were more likely to be stage 1 cancer (p < 0.0001). Subtyping was successful on 52 HPV31 samples. The HPV31 sublineages present were A1 (59%), A2 (4%), B2 (8%), and C3 (29%) (**Supplementary Table S5**). Due to the low number of A2 and B2 samples, statistics were only conducted on A1 and C3 samples. A1-positive samples were more likely to be cancer than C3 samples, as all 15 C3 samples were found in patients with precancer (p = 0.0014).

To investigate potential reasons no individuals with HPV31-C3 were found to have cancer, gene expression was compared between HPV31-C3 and the other HPV31 subtypes (**Supplementary Fig. 3**). Five genes were found to be differentially expressed. Four genes had higher expression in HPV31-C3: *CXCL14* (p = 0.040), *IRF-1* (p = 0.037), *HLA-B* (p = 0.044), and *HLA-E* (p = 0.0078). One gene, *CCNA2*, had higher expression in HPV31-A1, HPV31-A2, and HPV31-B2 (p = 0.012).

### 3.5 Gene expression during cancer progression

To understand differences in HPV and host gene expression during cervical cancer progression, we designed a panel of probes for gene expression of HPV16, HPV18, and HPV45 E6 and E7 gene expression as well as for 51 genes involved in cervical cancer or as markers for the tumor microenvironment (**Supplementary Table S1**). After quality control, 219 individuals and 50 genes remained (**Supplementary Table S6, Supplementary Fig. 1**).

Comparing control, precancer, and Stage 1 cancer showed 42 genes with significant differences before multiple test corrections, and selected genes are displayed in **Fig. 3**. The *APOBEC3A* and *APOBEC3B* anti-viral genes show increased expression in both precancer and cancer compared to controls (p < 0.0001, p < 0.0001). The expression of the cell proliferation promoter *MELTF* is higher in cancer compared to precancer and controls (p < 0.0001). Similarly, the cyclin genes *CCNA2* and *CCNE1* are higher in precancer and cancer than controls (p < 0.0001, p < 0.0001). CD28, forkhead box P3 (FOXP3), and indoleamine 2,3-dioxygenase 1 (IDO1) are markers of subsets of T cells. *CD28* expression is predominantly expressed by CD4+ T cells and is higher in controls than precancer or cancer (p < 0.0001). *FOXP3*, found on regulatory T cells, and *IDO1* (macrophages and dendritic cells) are higher in cancer than precancers or controls (p < 0.0001, p < 0.0001) (**Fig. 3E, F**). Poliovirus receptor (PVR), which can bind to multiple receptors to regulate the immune system and is known to promote tumor proliferation, is increased in cancer compared to precancer (p = 0.0003) [25]. FMS-related tyrosine kinase ligand (*FLT3LG*) is produced by NK cells and promotes dendritic cell proliferation [26]. This gene has lower expression in cancer than in precancers and controls (p = 0.0002) (**Fig. 3**). The expression of the NK cell receptor ligand *ULBP2* is increased in cancer compared to precancers and controls (p < 0.0001). Although *NECTIN2* expression differed between HPV16 and HPV31 precancers, there was no significant difference in *NECTIN2* expression between controls, precancer, and cancer. Unsurprisingly, the tumor suppressors tumor protein 53 (*TP53*) and retinoblastoma (*RB1*) have lower expression in cancer than precancer (p < 0.0001, p < 0.0001). The runt-related transcription factors 2 and 3 (*RUNX2* and *RUNX3*) genes are more highly expressed in controls and precancers than in cancers (p < 0.0001, p < 0.0001) (**Fig. 3G, H**). Multiple DNA repair genes are more highly expressed in cancer (**Supplementary Fig. 3**). Therefore, genes involved in innate and adaptive immunity, cell cycle regulation, and virally activated transcriptional regulation are differentially expressed during cervical cancer progression.

**Fig. 3.**
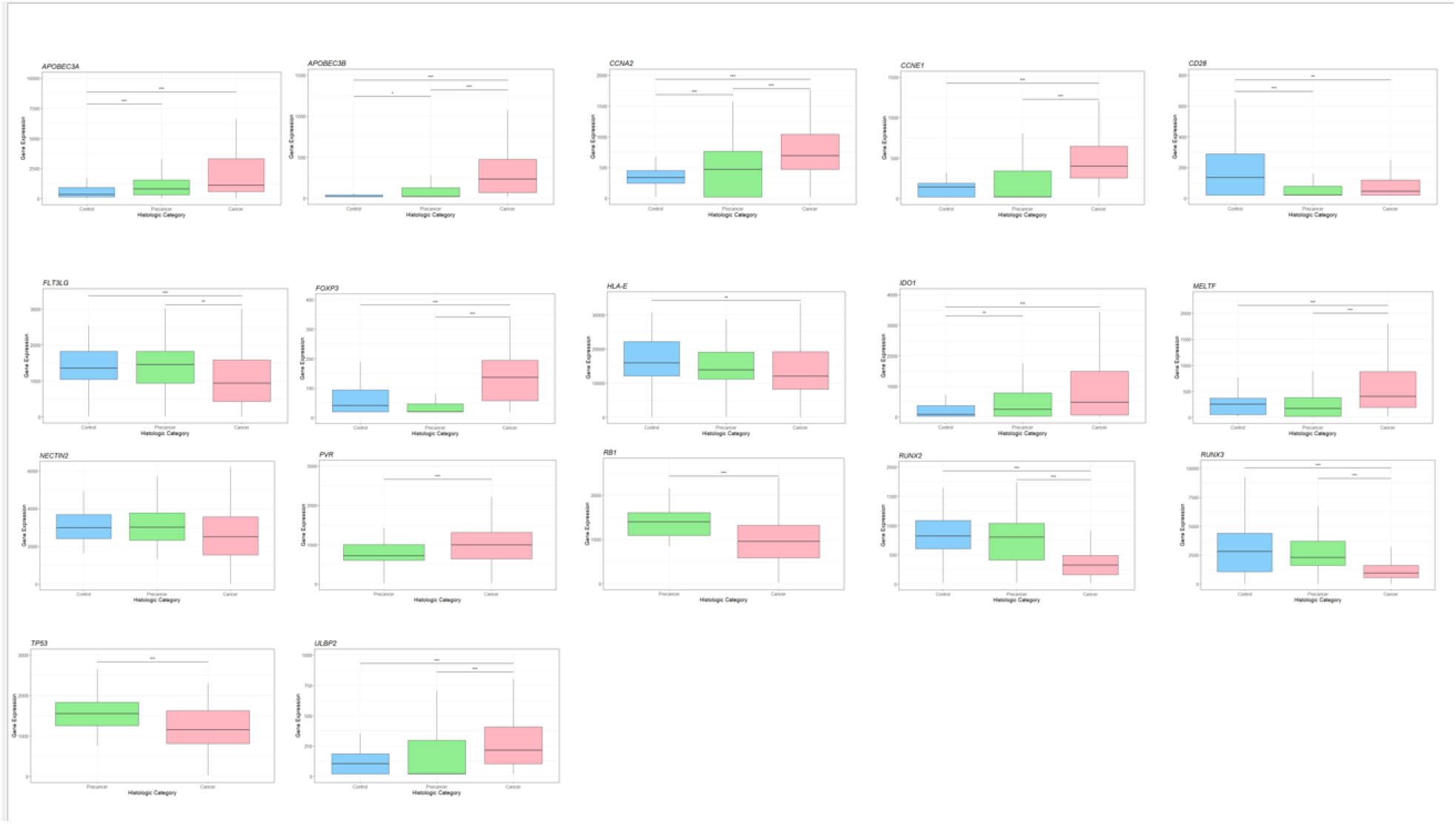
Analysis of individual gene expression in controls, precancer, and cancer. These box and whisker plots represent the expression level of various genes corresponding to their histological category, and gene expression is shown on the y-axis. In contrast, the histological type is shown on the x-axis. Expression of innate immune system genes (*APOBEC3A* and *APOBEC3B*), cell cycle genes (*CCNA2* and *CCNE1*), and immune cell markers (*CD28* and *FLT3LG*) is decreased in individuals with cancer. In contrast, expression of *FOXP3, IDO1*, and *ULBP2*, markers of immune system suppression and evasion, is increased. The expression of tumor suppressors (*TP53, RB1, RUNX2*, and *RUNX3*) is decreased in individuals with cancer. The expression of the NK-cell co-stimulator and co-inhibitor *NECTIN2* does not significantly change between controls, precancer, and cancer. *** indicates significance of p < 0.001, ** indicates significance of p < 0.01, and * indicates significance of p < 0.05.

### 3.6 Survival analyses of differentially expressed genes

For selected genes that differ significantly between controls, precancer, and cancer, we determined whether they could also predict survival in TCGA CESC (**Supplementary Table S7**). Genes associated with the innate and adaptive immune system were significantly associated with survival outcomes. The APOBEC3 proteins, markers of the innate immune system, are cytidine deaminases related to the cellular response to retroviruses, herpesviruses, and papillomaviruses [27]. Among individuals with all types of cervical cancer in the TCGA cohort, low *APOBEC3B* expression was significantly associated with worse survival (**Fig. 4**). Low *APOBEC3A* expression was associated with worse survival in individuals with squamous cell carcinoma (**Fig. 4**). These findings demonstrate the role of the APOBEC3 family in the immune response to HPV infection. APOBEC3 is stimulated upon infection by HPV, leading to higher expression in individuals with cancer and precancers. However, individuals with cancer who did not mount an immune response upon HPV infection then have poorer survival.

**Fig. 4.**
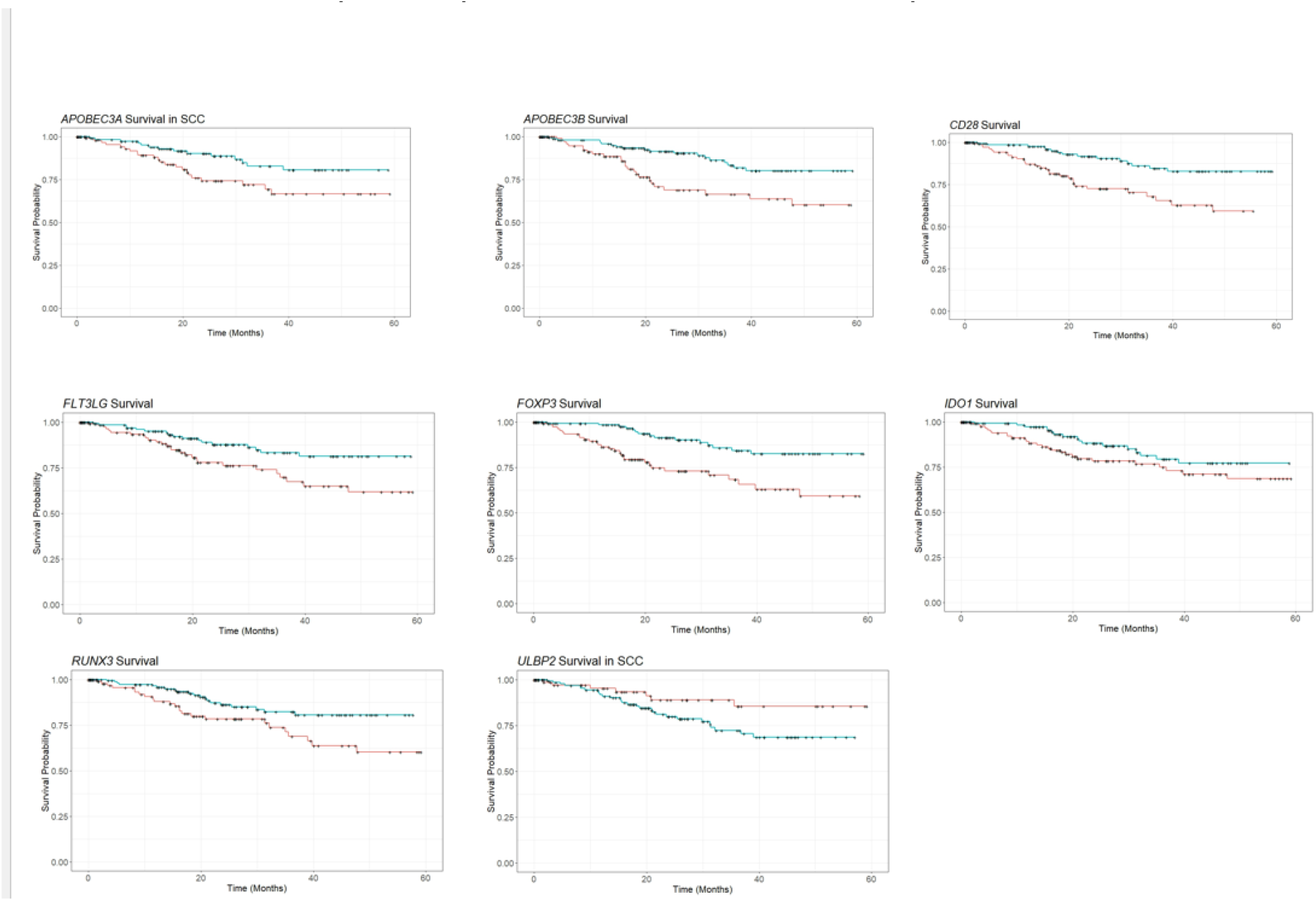
Survival of patients by expression of selected genes. These Kaplan-Meier curves show how expression levels of selected genes correspond with survival time. Time in months is shown on the x-axis, and the probability of survival is shown on the y-axis. The red line represents patients with lower expression, while the teal line represents patients with higher expression. Individuals with higher expression of *APOBEC3B, CD28, FLTLG3, FOXP3, IDO1*, and *RUNX3* had significantly better survival than individuals with lower expression. In patients with SCC, higher expression of *APOBEC3A* was associated with improved survival, while lower expression of *ULBP2* was associated with enhanced survival.

In the TCGA cohort, individuals with higher *CD28* and individuals with higher *FLT3LG* expression had higher survival than those with lower *CD28* expression and those with lower *FLT3LG* expression, showing that less immunosuppressed individuals were better able to combat cancer (**Fig. 4**) [28]. Interestingly, individuals with higher *FOXP3* and individuals with higher *IDO1* expression also had higher survival than those with lower expression of those genes. This finding was surprising, as high *FOXP3* expression and high *IDO1* expression indicate immunosuppression (**Fig. 4**). However, high *FOXP3* expression has been observed to correspond to higher survival in other cancers, such as gastric cancer, colorectal cancer, and non-small cell lung cancer [29-31]. High *IDO1* expression corresponds with higher survival in ovarian cancer [32]. In squamous cell carcinoma only, individuals with higher *ULBP2* expression had poorer survival, showing that cancer cells with stronger immune evading properties lead to a worse prognosis for patients (**Fig. 4**).

## 4. Discussion

Not much is known about the progression of cervical precancer to cancer in LMICs; therefore, we studied a cohort of women referred for colposcopy in Guatemala. We examined the HPV type distribution among precancer, stage 1, and invasive cancer samples. We found that HPV16, 18, and 45 have a significantly higher prevalence in invasive cancer, as has been documented in other populations [4]. Surprisingly, we found that HPV31 has a significantly higher prevalence in precancer in Guatemala (**Fig. 5**). We also performed gene expression quantitation in cervical cancer controls, precancer, and stage 1 cancer. Expression of innate immune system markers (*APOBEC3A/B*), DNA repair genes (*BRCA1, BRCA2, PALB2, RAD51*), cell cycle genes (*CCNA2* and *CCNE1*), a cell proliferation gene (*MELTF*), and a T-cell regulator (Treg) marker (*FOXP3*) was elevated in cancer compared to precancer and controls. Expression of *IDO1* was elevated in precancer and cancer as compared to controls. The expression of the tumor suppressor genes *RUNX2/3* decreased in cancer compared to precancer and controls. Additionally, the expression of the tumor suppressor genes *TP53* and *RB1* was decreased in cancer compared to precancer. The expression of *CD28* decreased in both cancer and precancer compared to the control group (**Fig. 5**), while the expression of *FLT3LG* was decreased in cancer compared to precancer and controls.

**Fig. 5.**
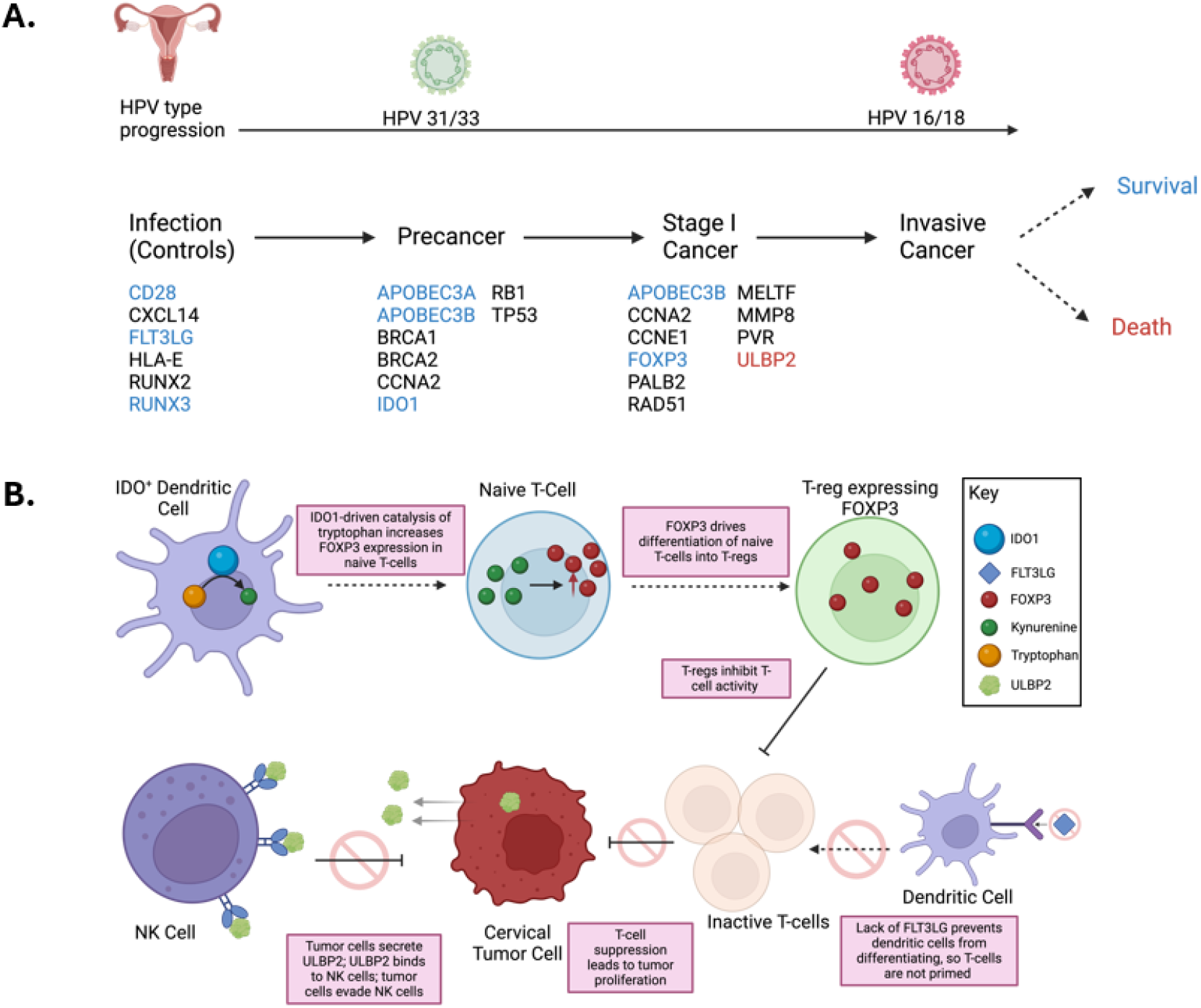
Changes in the expression of adaptive immune genes drive cervical cancer cell proliferation. A. HPV31 and HPV33 are more prevalent in precancer, while HPV16 and HPV18 are more prevalent in cancer. Genes listed under Infection (Controls) have significantly higher expression prior to the development of precancer. Genes listed under precancer increase in expression between controls and precancer cases. Genes listed under cancer increase in expression between precancer cases and cancer cases. Increased expression of genes listed in blue is associated with improved survival, while increased expression of genes listed in red is associated with worse survival. B. Increased expression of *IDO1* promotes *FOXP3* expression, which promotes proliferation of T-regulatory cells. Decreased expression of *FLT3LG* prevents the maturation of dendritic cells. T-cell suppression allows cervical tumor cells to proliferate. Expression of soluble *ULBP2* by cervical tumor cells inhibits NK cells, allowing immune evasion by tumor cells.

Worldwide, HPV16 is the most prevalent high-risk HPV type in invasive cervical cancer and causes nearly all HPV-positive head and neck, anal, and penile tumors[33]. The E6 oncoprotein of HPV16 has been shown to more strongly inactivate TP53 than other high-risk types[34]. In our Guatemalan sample, we observed a dramatically higher prevalence of HPV31 (32%) in cervical precancer samples and a much lower prevalence in invasive cancer (4%). In comparison, the prevalence of HPV16 was much lower in individuals with precancer (23%) than in invasive cancer (47%). Individuals with precancer had 6.38 (95% CI 3.15, 12.77) times the odds of having HPV31 than individuals with stage one and invasive cancer, and individuals with invasive cancer had 2.39 (95% CI 1.61, 3.60) times the odds of having HPV16 than individuals with precancer and stage one cancer. Based on these odds ratios, we compared gene expression between HPV16 and HPV31 to elucidate the significant genes driving the transition from precancer to cancer.

We first examined three genes that regulate cell proliferation. *CCNE1* encodes the cyclin E1 proteins and is crucial in cell cycle regulation. CCNE1 partners with cyclin-depen dent kinases (CDKs) to drive the transition from the G1 to S phase. [35, 36]. The expression of *MELTF*, which encodes the protein melanotransferrin, increases in several cancers, including lung adenocarcinoma, ovarian cancer, and melanoma. The molecular mechanisms of MELTF are not fully understood, but previous research suggests that it regulates cell proliferation. In melanoma, downregulation of *MELTF* inhibits tumor proliferation[17]. In ovarian cancer, *MELTF* drives the epithelial to mesenchymal transition, leading to metastasis[15]. In lung cancer, *MELTF* promotes tumor growth by stimulating the Notch pathway. Our analysis showed increased *CCNE1* and *MELTF* expression in cancer. In precancer, expression of the two genes is increased in HPV16 compared to HPV31. The increased stimulation of CCNE1 and *MELTF* by HPV16 could explain why individuals with HPV16-positive precancer are more likely to progress to cancer than individuals with HPV31-positive precancer.

The APOBEC3 antiviral response is important in HPV control [37]. We identified a significant increase in *APOBEC3A/B* expression in cancer compared to precancers and controls. Increased expression of *APOBEC3B* in individuals with cancer is associated with higher survival in TCGA, indicating that a stronger response to viral infection is correlated with a better prognosis.

The adaptive immune response eliminates HPV-infected cells and precancerous lesions [38]. Our study indicates that the adaptive immune system is heavily suppressed in cervical cancer patients by inhibiting T-cell proliferation. *FLT3LG* promotes the differentiation of dendritic cells, which prime T-cells [39]. We found that individuals with cancer had lower levels of *FLT3LG* expression, indicating decreased activity of dendritic cells and T-cells. Similarly, we found increased expression of Indoleamine-2,3-dioxygenase 1 (*IDO1*), which catabolizes tryptophan to generate immunosuppressive kynurenines [40]. Kynurenines regulate T-cell differentiation by inducing the expression of the transcription factor *FOXP3 [41]*. Expression of FOXP3 causes naive CD4+ cells to become T-regulatory cells, leading to immunosuppression. FOXP3 is, therefore, a marker of T-regulatory cells. Both FOXP3 and IDO1 had increased expression in cancer, further indicating T-cell suppression. These findings are supported by our finding of decreased expression of *CD28* in cancer, as CD28 is expressed on both resting and activated T-cells [42, 43].

To extend our findings, we examined the association between markers of the adaptive immune system and patient survival in TCGA cervical cancer. The expression of *CD28* and *FOXP3* was highly correlated, and high expression of *IDO1, FOXP3, CD28*, and *FOXP3* was all associated with improved survival in cervical cancer. Increased *CD28* expression has been shown to correlate with improved survival in head and neck squamous cell carcinoma (HNSCC), while increased *FLT3LG* expression has previously been shown to correlate with improved survival in cervical cancer [28, 44]. While the improved survival of patients with increased expression of *FOXP3* or *IDO1* contradicts the role of T-cell suppression in cervical cancer, *FOXP3* and *IDO1* can be expressed in tumor cells themselves [45, 46]. Previous studies have shown a positive association between nuclear *FOXP3* expression in tumor cells and survival in breast cancer, gastric cancer, and hepatocellular carcinoma [29, 45, 47]. Similarly, in ovarian cancer, increased expression of *IDO1* is associated with improved prognosis[32]. Follow-up studies are needed to examine the expression of *FOXP3* and *IDO1* in cervical tumor cells and infiltrating T-cells.

In addition to the suppression of T-cells, our data on the expression of *ULBP2* and *HLA-E* shows that inhibition of NK cells also drives progression from precancer to cancer. *ULBP2* is a ligand of the natural killer group 2, member D (NKG2D) activating receptor, expressed on NK cells. NKG2D binds to stress-related ligands expressed on tumor cells to initiate the antitumor immune response [48]. Although membrane-bound *ULBP2* drives tumor immunity, some tumors can secrete soluble ULBP2 to evade NK-cell-driven immunosurveillance [18]. Soluble ULBP2 decreases the expression of membrane-bound ULBP2 and inhibits NKG2D [18]. Our analysis showed a higher expression of *ULBP2* in individuals with cancer, and increased expression of *ULPB2* was correlated with poorer survival, indicating that *ULBP2* acts as an NK-cell inhibitor in cervical cancer. We also observed that *ULBP2* expression is increased in HPV16-positive precancers as compared to HPV31-positive precancers, indicating that HPV16-positive precancers are more robust at evading immunosurveillance and, therefore more likely to progress to cancer than HPV31-positive precancers. HLA-E interacts with the NK-cell receptor *NKG2A* to promote the education of NK cells. Our analysis showed a decreased expression of *HLA-E* in individuals with cancer, demonstrating an inhibited immune response. We also observed a reduced expression in individuals with HPV16-positive precancers as compared to individuals with HPV31-positive precancers, further supporting enhanced immune evasion by HPV16-positive precancers.

HPV does not encode enzymes for DNA replication and hijacks host cells’ mechanisms to proliferate. HPV infection leads to abnormal expression of DNA repair enzymes, contributing to genome instability. Expression of the DNA repair enzymes *BRCA1/2, PALB2*, and *RAD51* were increased in cancer samples compared to controls, indicating that double-stranded DNA repair activity increases during cervical cancer progression. Expression of *PALB2* was increased in HPV16-positive precancers as compared to HPV31-positive precancers, further supporting its role in cervical cancer progression.

The RUNX gene family plays a role in cancer development and progression and is a tumor suppressor or promoter, depending on the tumor type. Previous studies have shown that *RUNX3* is a tumor suppressor in cervical cancer [49], we found that expression of *RUNX2* and *RUNX3* was decreased in cervical cancer cells compared to controls. Furthermore, in TCGA, increased expression of *RUNX3* was associated with better survival. RUNX genes also regulate the activation of immune cells in the tumor microenvironment [50] and can have a complex on anti-tumor immunity.

Our study had limited demographic data and no follow-up or survival data. In our analysis of HPV progression to cancer, our sample size was insufficient to accurately determine the rate of progression of individual HPV types. We have not determined the genetic ancestry of the subjects and cannot assess the impact of European versus Indigenous American backgrounds. The gene panel was limited and did not include all genes relevant to cancer progression. Many samples were excluded due to low gene expression, possibly introducing bias.

Several immune therapies, including checkpoint inhibitors and engineered T-cells, have proven effective in treating cervical cancer [51, 52]. Our findings suggest that further studies of immune cells in the tumor microenvironments of cervical precancer and cancer could aid the development of therapeutic approaches. While germline mutations in homologous recombination repair genes such as *BRCA1/2* are not associated with cervical cancer, our data suggests that targeted therapies to this pathway could be beneficial. Several CDK inhibitor drugs are currently being explored in cancer treatments[53]. Our finding that regulators of CDKs are elevated in cervical cancer suggests that these drugs could prove to be an effective form of treatment. LMIC countries have the highest incidence and case fatality rates of cervical cancer. As shown in this study, exploring a cohort of women in an LMIC can lead to insight into which biological factors influence the progression of cervical cancer. Building on the knowledge of these biological changes, as well as the current immunotherapies being developed can establish a clear pathway from diagnosis to treatment.

## Conclusions

To better understand the progression from HPV infection to cervical precancer and cancer, we have analyzed a set of biopsy samples from women referred to colposcopy from a single center in Guatemala. Analysis of HPV types documented differential risk progression of selected HPV types, and we had a sufficient sample size of HPV16 (high progression risk) and HPV31 (low progression risk) to examine differences in gene expression of HPV16 and 31 precancers. We identified genes involved in proliferation (*CCNE1, MELTF*) and immune escape (*ULBP2, HLA*-E, *CD28*). In addition, we identified higher expression of cell cycle genes (*CCNA2* and *CCNE1*) in precancers compared to disease-free controls and higher expression of immune genes (*CD28, FOXP3*, and *IDO1*) in controls. Together, these findings provide insight into the progression of cervical cancer in an understudied population.

## Data Statement

The data from the manuscript is available upon request and will be deposited in the xxx repository.

## Author contributions: CRediT

Conceptualization MD, RO; Formal analysis, ER, LB, SKM, MY; Methodology IR, HJL. LB, VA, YX, HL, RM, HJL; Writing – original draft, ER, IR, MD; Writing – review and editing VR, YX, HL, LM.

## Acknowledgements

We thank Adolfo Santizo and Lineth Boror for technical assistance in sample collection.

## Abbreviations

Treg: T-regulatory cell
LMIC: Low and middle-income countries
hrHPV: High-risk HPVs
SCC: Squamous Cell Carcinoma
NK cells: Natural Killer cells
FFPE: Formalin-fixed paraffin-embedded
DSCF: Dwass-Steel-Critchlow-Fligner
TCGA: The Cancer Genome Atlas
NCI CHANGeS: National Cancer Institute Carcinogenic HPV All Next Generation Sequencing
CDK: Cyclin Dependant Kinases
INCan: Instituto Nacional de Cancerologia

## References

1. Schiffman, M., et al., Carcinogenic human papillomavirus infection. Nat Rev Dis Primers, 2016. 2: p. 16086.

2. Rodriguez, A.C., et al., Rapid clearance of human papillomavirus and implications for clinical focus on persistent infections. J Natl Cancer Inst, 2008. 100(7): p. 513–7.

3. Demarco, M., et al., A study of type-specific HPV natural history and implications for contemporary cervical cancer screening programs. EClinicalMedicine, 2020. 22: p. 100293.

4. Guan, P., et al., Human papillomavirus types in 115,789 HPV-positive women: a meta-analysis from cervical infection to cancer. Int J Cancer, 2012. 131(10): p. 2349–59.

5. Singh, D., et al., Global estimates of incidence and mortality of cervical cancer in 2020: a baseline analysis of the WHO Global Cervical Cancer Elimination Initiative. Lancet Glob Health, 2023. 11(2): p. e197–e206.

6. Sung, H., et al., Global Cancer Statistics 2020: GLOBOCAN Estimates of Incidence and Mortality Worldwide for 36 Cancers in 185 Countries. CA Cancer J Clin, 2021. 71(3): p. 209–249.

7. den Boon, J.A., et al., Molecular transitions from papillomavirus infection to cervical precancer and cancer: Role of stromal estrogen receptor signaling. Proc Natl Acad Sci U S A, 2015. 112(25): p. E3255–64.

8. Zhai, Y., et al., Gene expression analysis of preinvasive and invasive cervical squamous cell carcinomas identifies HOXC10 as a key mediator of invasion. Cancer Res, 2007. 67(21): p. 10163–72.

9. Lou, H., et al., Genome Analysis of Latin American Cervical Cancer: Frequent Activation of the PIK3CA Pathway. Clin Cancer Res, 2015. 21(23): p. 5360–70.

10. Rossi, N.M., et al., Extrachromosomal Amplification of Human Papillomavirus Episomes is a Mechanism of Cervical Carcinogenesis. Cancer Res, 2023.

11. Cullen, M., et al., Deep sequencing of HPV16 genomes: A new high-throughput tool for exploring the carcinogenicity and natural history of HPV16 infection. Papillomavirus Res, 2015. 1: p. 3–11.

12. Untergasser, A., et al., Primer3--new capabilities and interfaces. Nucleic Acids Res, 2012. 40(15): p. e115.

13. Cancer Genome Atlas Research, N., et al., Integrated genomic and molecular characterization of cervical cancer. Nature, 2017. 543(7645): p. 378–384.

14. Gao, J., et al., Integrative analysis of complex cancer genomics and clinical profiles using the cBioPortal. Sci Signal, 2013. 6(269): p. pl1.

15. Ji, L., et al., Knockout of MTF1 Inhibits the Epithelial to Mesenchymal Transition in Ovarian Cancer Cells. J Cancer, 2018. 9(24): p. 4578–4585.

16. Zhang, L. and L. Shi, The E2F1/MELTF axis fosters the progression of lung adenocarcinoma by regulating the Notch signaling pathway. Mutat Res, 2023. 827: p. 111837.

17. Dunn, L.L., E.O. Sekyere, Y. Suryo Rahmanto, and D.R. Richardson, The function of melanotransferrin: a role in melanoma cell proliferation and tumorigenesis. Carcinogenesis, 2006. 27(11): p. 2157–69.

18. Waldhauer, I. and A. Steinle, Proteolytic release of soluble UL16-binding protein 2 from tumor cells. Cancer Res, 2006. 66(5): p. 2520–6.

19. Zhu, Y., et al., Identification of CD112R as a novel checkpoint for human T cells. J Exp Med, 2016. 213(2): p. 167–76.

20. Lin, Z., et al., HLA class I signal peptide polymorphism determines the level of CD94/NKG2-HLA-E-mediated regulation of effector cell responses. Nat Immunol, 2023. 24(7): p. 1087–1097.

21. Townsend, S.E. and J.P. Allison, Tumor rejection after direct costimulation of CD8+ T cells by B7-transfected melanoma cells. Science, 1993. 259(5093): p. 368–70.

22. Perevalova, A.M., L.F. Gulyaeva, and V.O. Pustylnyak, Roles of Interferon Regulatory Factor 1 in Tumor Progression and Regression: Two Sides of a Coin. Int J Mol Sci, 2024. 25(4).

23. Mirabello, L., et al., HPV16 Sublineage Associations With Histology-Specific Cancer Risk Using HPV Whole-Genome Sequences in 3200 Women. J Natl Cancer Inst, 2016. 108(9).

24. Pinheiro, M., et al., Phylogenomic Analysis of Human Papillomavirus Type 31 and Cervical Carcinogenesis: A Study of 2093 Viral Genomes. Viruses, 2021. 13(10).

25. Liu, W.F., et al., PVR-A Prognostic Biomarker Correlated with Immune Cell Infiltration in Hepatocellular Carcinoma. Diagnostics (Basel), 2022. 12(12).

26. Barry, K.C., et al., A natural killer-dendritic cell axis defines checkpoint therapy-responsive tumor microenvironments. Nat Med, 2018. 24(8): p. 1178–1191.

27. Stavrou, S. and S.R. Ross, APOBEC3 Proteins in Viral Immunity. J Immunol, 2015. 195(10): p. 4565–70.

28. Chen, L., et al., Low Level FLT3LG is a Novel Poor Prognostic Biomarker for Cervical Cancer with Immune Infiltration. J Inflamm Res, 2022. 15: p. 5889–5904.

29. Ma, G.F., et al., High FoxP3 expression in tumour cells predicts better survival in gastric cancer and its role in tumour microenvironment. Br J Cancer, 2014. 110(6): p. 1552–60.

30. Sun, X., et al., Expression of Foxp3 and its prognostic significance in colorectal cancer. Int J Immunopathol Pharmacol, 2017. 30(2): p. 201–206.

31. Yang, S., et al., FOXP3 promotes tumor growth and metastasis by activating Wnt/betacatenin signaling pathway and EMT in non-small cell lung cancer. Mol Cancer, 2017. 16(1): p. 124.

32. Hoffmann, I., et al., Increased expression of IDO1 is associated with improved survival and increased number of TILs in patients with high-grade serous ovarian cancer. Neoplasia, 2023. 44: p. 100934.

33. Olesen, T.B., et al., Prevalence of human papillomavirus DNA and p16(INK4a) in penile cancer and penile intraepithelial neoplasia: a systematic review and meta-analysis. Lancet Oncol, 2019. 20(1): p. 145–158.

34. Mesplede, T., et al., p53 degradation activity, expression, and subcellular localization of E6 proteins from 29 human papillomavirus genotypes. J Virol, 2012. 86(1): p. 94–107.

35. Yao, S., et al., Clinical characteristics and outcomes of phase I cancer patients with CCNE1 amplification: MD Anderson experiences. Sci Rep, 2022. 12(1): p. 8701.

36. Jiang, A., et al., CCNA2 as an Immunological Biomarker Encompassing Tumor Microenvironment and Therapeutic Response in Multiple Cancer Types. Oxid Med Cell Longev, 2022. 2022: p. 5910575.

37. Warren, C.J., J.A. Westrich, K.V. Doorslaer, and D. Pyeon, Roles of APOBEC3A and APOBEC3B in Human Papillomavirus Infection and Disease Progression. Viruses, 2017. 9(8).

38. Litwin, T.R., et al., Infiltrating T-cell markers in cervical carcinogenesis: a systematic review and meta-analysis. Br J Cancer, 2021. 124(4): p. 831–841.

39. Cueto, F.J. and D. Sancho, The Flt3L/Flt3 Axis in Dendritic Cell Biology and Cancer Immunotherapy. Cancers (Basel), 2021. 13(7).

40. Pallotta, M.T., et al., Indoleamine 2,3-dioxygenase 1 (IDO1): an up-to-date overview of an eclectic immunoregulatory enzyme. FEBS J, 2022. 289(20): p. 6099–6118.

41. Stone, T.W. and R.O. Williams, Interactions of IDO and the Kynurenine Pathway with Cell Transduction Systems and Metabolism at the Inflammation-Cancer Interface. Cancers (Basel), 2023. 15(11).

42. Tai, X., M. Cowan, L. Feigenbaum, and A. Singer, CD28 costimulation of developing thymocytes induces Foxp3 expression and regulatory T cell differentiation independently of interleukin 2. Nat Immunol, 2005. 6(2): p. 152–62.

43. Soligo, M., et al., CD28 costimulation regulates FOXP3 in a RelA/NF-kappaB-dependent mechanism. Eur J Immunol, 2011. 41(2): p. 503–13.

44. Chang, S.R., et al., The Concordant Disruption of B7/CD28 Immune Regulators Predicts the Prognosis of Oral Carcinomas. Int J Mol Sci, 2023. 24(6).

45. Takenaka, M., et al., FOXP3 expression in tumor cells and tumor-infiltrating lymphocytes is associated with breast cancer prognosis. Mol Clin Oncol, 2013. 1(4): p. 625–632.

46. Liu, M., et al., Targeting the IDO1 pathway in cancer: from bench to bedside. J Hematol Oncol, 2018. 11(1): p. 100.

47. Gong, Z., et al., Nuclear FOXP3 inhibits tumor growth and induced apoptosis in hepatocellular carcinoma by targeting c-Myc. Oncogenesis, 2020. 9(10): p. 97.

48. Liu, H., et al., Role of NKG2D and its ligands in cancer immunotherapy. Am J Cancer Res, 2019. 9(10): p. 2064–2078.

49. Li, Z., P. Fan, M. Deng, and C. Zeng, The roles of RUNX3 in cervical cancer cells in vitro. Oncol Lett, 2018. 15(6): p. 8729–8734.

50. Ito, Y., S.C. Bae, and L.S. Chuang, The RUNX family: developmental regulators in cancer. Nat Rev Cancer, 2015. 15(2): p. 81–95.

51. Nagarsheth, N.B., et al., TCR-engineered T cells targeting E7 for patients with metastatic HPV-associated epithelial cancers. Nat Med, 2021.

52. Stevanovic, S., et al., A Phase II Study of Tumor-infiltrating Lymphocyte Therapy for Human Papillomavirus-associated Epithelial Cancers. Clin Cancer Res, 2019. 25(5): p. 1486–1493.

53. Goel, S., J.S. Bergholz, and J.J. Zhao, Targeting CDK4 and CDK6 in cancer. Nat Rev Cancer, 2022. 22(6): p. 356–372.

