## Supplemental Figures for "Analysis of the progression of cervical cancer in Guatemala- from pre-malignancy to invasive disease"

1. **S1. Diagram/flow chart of what happened to all the samples**
2. **S2 –insert S3 HPV31 sublineage figures here probably–**
3. **S3 DNA repair genes**

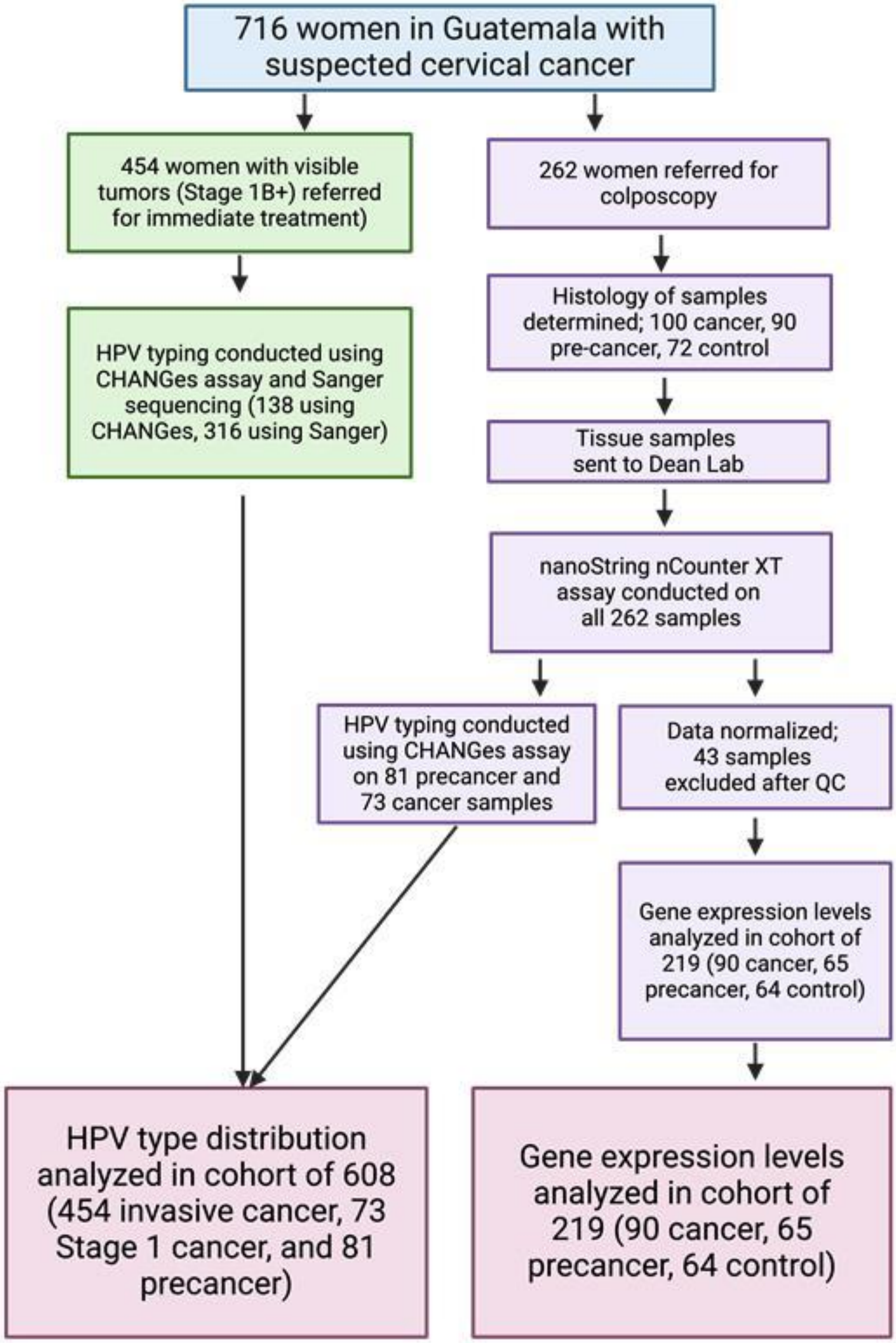


**Figure S1. Sample flowthrough and exclusions.**

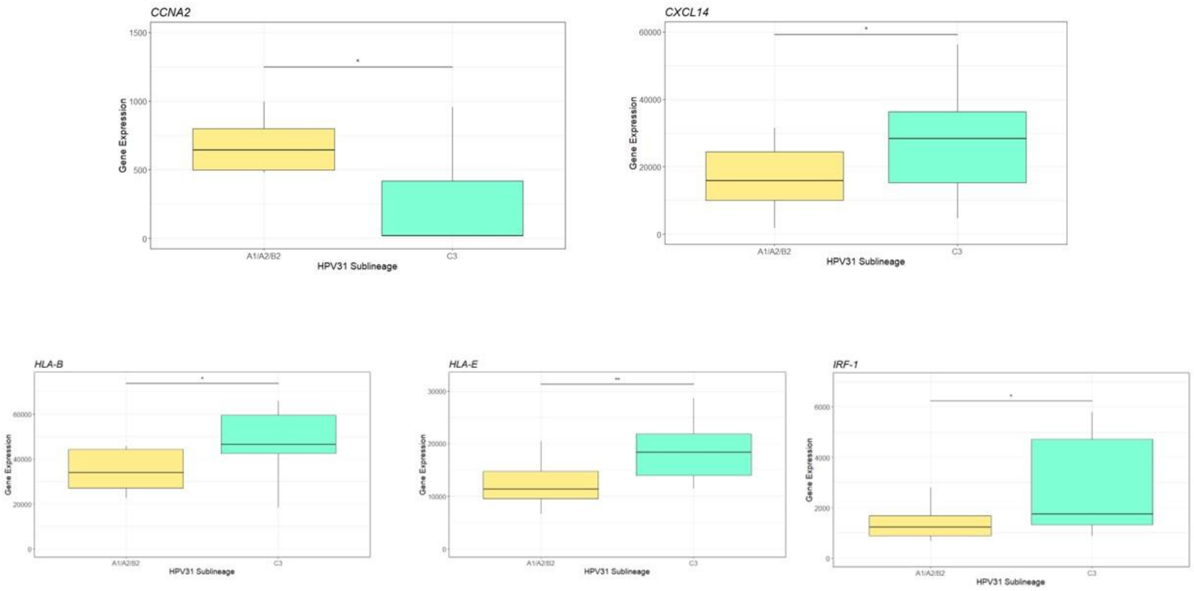

**Supplementary Fig. S2. Differences in gene expression between HPV31-C3-positive precancers and HPV31-positive precancers of other sublineages.** Box and whisker plots representing the expression level of genes differentially expressed in HPV31-C3-positive precancers and HPV31-positive precancers of other sublineages. Gene expression is shown on the y-axis, while HPV type is on the x-axis. The expression of *CCNA2* is lower in HPV31-C3-positive precancers, while the expression of *CXCL14*, *IRF-1*, *HLA-B*, and *HLA-E* is higher in HPV31-C3-positive precancers.

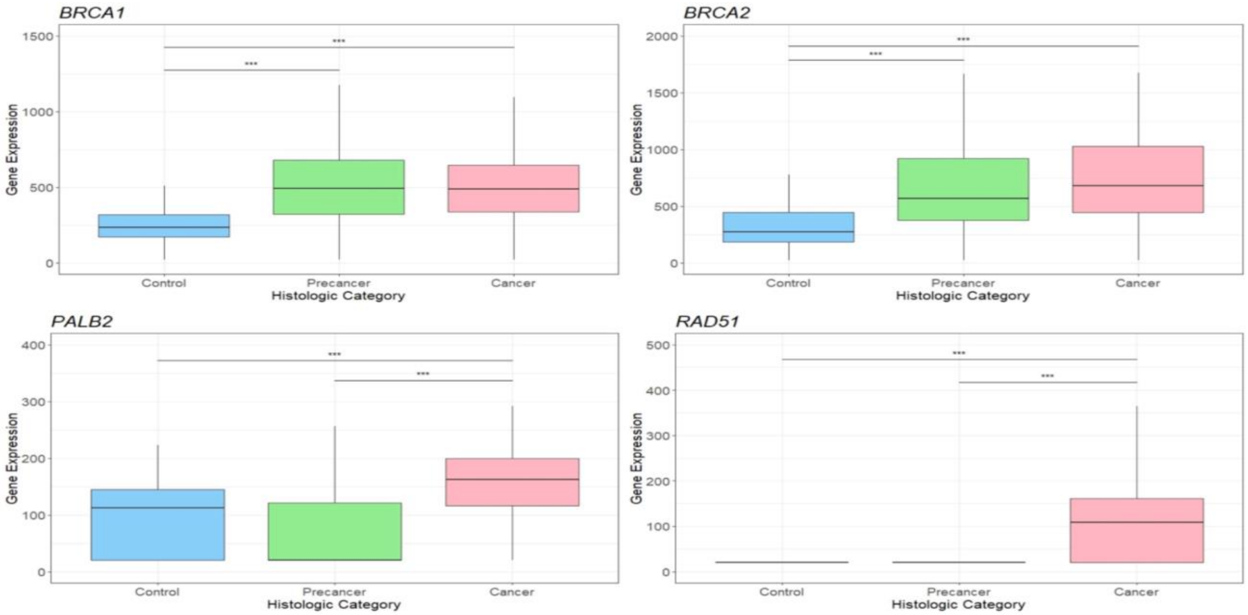


**Supplementary Fig. S3. Analysis of DNA repair gene expression in controls, precancer, and cancer.** These box and whisker plots represent the expression level of DNA repair genes corresponding to their histological category. Gene expression is shown on the y-axis, while the histological type is shown on the x-axis. The expression of *BRCA1* and *BRCA2* increases in cancer and precancers compared to the control group. The expression of *PALB2* and *RAD51* increases in cancer compared to precancer and controls.  *** indicates significance of $p < 0.001$, ** indicates significance of $p < 0.01$, and * indicates significance of $p < 0.05$.
